# Identification of *SENP7* and *UTF1*/*VENTX* as new loci influencing clustered protocadherin methylation across blood and brain using a genome-wide association study

**DOI:** 10.1101/2024.12.19.629385

**Authors:** Yunfeng Liu, Maja Vukic, Eilis Hannon, Hailiang Mei, BIOS Consortium, Jonathan Mill, Lucia Daxinger, Bastiaan T. Heijmans

## Abstract

The epigenetic regulation of clustered protocadherin (c*PCDH*) genes is tightly linked to their function as specific cell surface barcodes for neural self-nonself discrimination. Differential c*PCDH* DNA methylation in both blood and brain has been implicated in diverse human traits and diseases ranging from BMI, aging, cancer and brain disorders. However, the unique regulation of c*PCDH* methylation remains poorly understood. Therefore, we performed a genome-wide association study to evaluate the association of >7 million genetic variants with DNA methylation at 607 c*PCDH* CpGs measured in whole blood of 3777 individuals and validated findings in prefrontal cortex samples obtained from 523 brain donors. We observed concordant c*PCDH* methylation patterns in blood and prefrontal cortex, which switched between hypo-, intermediate and hypermethylation over short distances with the former overlapping with the promoter regions of each c*PCDH* member. Through methylation quantitative trait locus (meQTL) analysis *in trans*, we first confirmed the broad effect of the candidate gene *SMCHD1* on c*PCDH* methylation in blood which validated in prefrontal cortex. Through a genome-wide analysis, we next identified the *SENP7* and *UTF1*/*VENTX* loci to have widespread, subcluster-specific effects on c*PCDH* methylation in blood which was validated in prefrontal cortex. While *SENP7* can indirectly affect DNA methylation through the deSUMOylation of the chromatin repressor *KAP1*, *UTF1* and *VENTX* are two genes involved in embryonic development not previously implicated in epigenetic regulation. Our findings shed new light on the processes involved in c*PCDH* methylation that may underlie associations with disease.

## Introduction

A broad range of human diseases and traits have been linked to changes in epigenetic regulation of the clustered protocadherin genes (c*PCDH*, mapping to chromosome 5q31)^1^. Epigenome-wide association studies reported differential methylation for BMI^2^, brain disorders^3–6^ and age^7,8^, while c*PCDH* also shows DNA methylation differences in multiple forms of cancer^9–11^. In addition, aberrant c*PCDH* methylation is characteristic for patients as affected by the immunodeficiency, centromeric region instability, facial anomalies (ICF) syndrome^12^ and facioscapulohumeral muscular dystrophy (FSHD)^13^. These epigenetic associations resulted from the study of various cell types including blood and brain, although c*PCDH* expression is predominantly observed in brain. Contrasting these well-described disease associations is the still limited insight in the regulation of c*PCDH* methylation.

In humans, the more than 50 clustered *PCDHA*, *PCDHB* and *PCDHG* genes together form a 1Mb locus, referred to as c*PCDH* genes. In the brain, the expression of the c*PCDH* genes is controlled by a unique combination of regulatory mechanisms. These include CCTC binding factor (CTCF)/cohesion mediated chromatin looping^14,15^, histone modifications^16,17^ and DNA methylation^18^. In each neuronal cell, the exact tuning of these mechanisms results in the stochastic expression of a different subset of c*PCDH* gene members^19^. The resulting combination serves as specific cell surface barcode that help neurons distinguish between self and nonself^20^. DNA methylation of c*PCDH* gene promoters play a key role in the stochastic expression of c*PCDH* members^18^. Specifically, methylation of the CpG dinucleotide at c*PCDH* promoters precludes CTCF binding, thus disrupting the formation of long-range chromatin loops^14^. The regulation of c*PCDH* methylation is only partly understood in humans. Studies in mice, however, identified various genes involved. Two well-described examples are *DNMT3B* and *SMCHD1*. *DNMT3B* regulates the stochastic expression of c*PCDH* members through mediating differential methylation of promoters in neurons^18^, while *SMCHD1* is required for c*PCDH* methylation and its loss induces alterations in c*PCDH* expression^21,22^. Two more recently discovered include the *WAPL* gene^23^, which functions as a rheostat of c*PCDH* member diversity, and we reported the role of the *CDCA7* gene^24^, which regulates c*PCDH* methylation and expression frequency. Human studies are necessary to shed light on the origin of differential c*PCDH* methylation implicated in human traits and diseases and may also indicate additional mechanisms underlying the generation of cell surface barcodes that is unique to c*PCDH* genes.

To map loci involved in the epigenetic regulation of c*PCDH* genes in humans, we performed a genome-wide analysis to discover common genetic variants that are associated with c*PCDH* methylation in blood (n=3777) followed by validation of these associations in prefrontal cortex (n=523). We not only found that the candidate gene *SMCHD1 locus* has widespread effects on c*PCDH* methylation in blood and brain, but also report two new loci to be involved, namely *SENP7*, a gene involved in deSUMOylation^25^, and the *UTF1*/*VENTX* locus, two genes involved in embryonic development^26,27^.

## Methods

### Discovery Cohorts

The discovery cohorts in this study was based on the Biobank-based Integrative Omics Studies (BIOS) Consortium, which comprises six Dutch biobanks: Cohort on Diabetes and Atherosclerosis Maastricht (CODAM)^28^, LifeLines (LL)^29^, Leiden Longevity Study (LLS)^30^, Netherlands Twin Register (NTR)^31^, Rotterdam Study (RS)^32^, and the Prospective ALS Study Netherlands (PAN)^33^. The data we analyzed consists of 3777 unrelated individuals for whom genotype data and DNA methylation data were available, and for 3013 individuals, RNA-seq data were available (**Supplementary Data 1, Table S1**). Genotype data, DNA methylation data and gene expression data were all measured in whole blood. The Human Genotyping facility (HuGe-F, Erasmus MC, Rotterdam, The Netherlands, http://www.glimdna.org) generated the methylation and RNA-sequencing data.

Genotype data were measured per cohort individually. Details on the data generation can be found in the individual papers—CODAM^34^, LL^29^, LLS^35^, NTR^36^, RS^32^, and PAN^33^. The genotype data were harmonized towards the Genome of the Netherlands (GoNL) using Genotype Harmonizer^37^ and subsequently imputed per cohort using MaCH^38^ with the Haplotype Reference Consortium (HRCv1.1) panel^39^. For each cohort, SNPs with R^2^ < 0.3 and call rate < 0.95 were removed, and then VCFtools^40^ was used to remove SNPs with Hardy-Weinberg equilibrium P-value < 10^−4^. After merging the cohorts, SNPs with minor allele frequency < 0.01 were removed. These imputation and filtering steps resulted in 7,751,736 SNPs that passed quality control in each of the datasets.

DNA methylation data of whole blood samples were obtained from bisulfite-converting 500 ng of genomic DNA using the Zymo EZ DNA Methylation kit (Zymo Research, Irvine, CA, USA). Next, 4 μl of bisulfite-converted DNA was hybridized on the Illumina HumanMethylation450 array according to the manufacturer’s protocol (Illumina, San Diego, CA, USA). Preprocessing and normalization of the data were done as described in DNAmArray workflow previously developed by our group (https://molepi.github.io/DNAmArray_workflow/). Briefly, original IDAT files were imported into by R package *minfi*^41^, followed by sample-level quality control (QC) was performed using *MethylAid*^42^. Filtering of probes was based on detection P value (*P* < 0.01), number of beads available (≤ 2), or zero values for signal intensity. Normalization was done using functional normalization as implemented in *minfi*^41^, using five principal components extracted using the control probes for normalization. All samples or probes with more than 5% missing were excluded. In addition, probes located on sex chromosomes, or that overlap with known common genetic variants or ambiguously mapping or cross-reactive recommended by Zhou et al were all removed for the reliability of analysis^43^. After these filtering steps, our final dataset consisted of 412,373 probes measured in 3777 samples. To exclude a negative impact of non-normal distributions and outliers on the validity of our results, DNA methylation data were transformed by rank-inverse normal (RIN) transformation for each cohort^44–46^.

Detailed information on the generation and processing of the RNA-seq data can be found in previous work^47^. In short, globin transcripts were removed from whole blood RNA using the Ambion GLOBINclear kit and subsequently processed for RNA-sequencing using the Illumina TruSeq version 2 library preparation kit. RNA libraries were paired-end sequenced using Illumina’s HiSeq 2000 platform with a read length of 2 × 50 bp, pooling 10 samples per lane. Reads which passed the chastity filter were extracted with CASAVA. Quality control was done in three steps: initial QC was performed using FastQC (v0.10.1), adaptor sequences were removed using Cutadapt, and read ends with insufficient quality were removed with Sickle. Reads were aligned to the human genome (hg19) using STAR (v2.3.0e). To avoid reference mapping bias, all GoNL SNPs (https://www.nlgenome.nl/) with MAF > 0.01 in the reference genome were masked with N. Read pairs with at most 8 mismatches, mapping to at most 5 positions, were used. Gene counts were calculated by summing the total number of reads aligning to a gene’s exons according to Ensembl, version 71. Samples for which less than 70% of all reads mapped to exons were removed. For data analysis, we used log counts per million (CPM), and only used protein coding genes with sufficient expression levels (median CPM > 1), resulting in 11,769 genes. Similar to DNA methylation data, RNA-seq data were transformed by rank inverse normal (RIN) transformation for each cohort^44–46^. Finally, correct data linkage of genotype, DNA methylation and RNA-seq data was verified using OmicsPrint^48^.

### Validation cohorts

Data generated on tissue from the Brains for Dementia Research (BDR) cohort, which consists of a network of six dementia research centers across England and Wales (based at Bristol, Cardiff, King’s College London, Manchester, Oxford, and Newcastle Universities) and five brain banks (brain donations from Cardiff are banked at King’s College London)^49^, was used to validate our results. Briefly, participants >65 years of age were recruited using both national and local press, TV and radio coverage as well as at memory clinics and support groups. The cohort includes those with and without dementia and covers the full range of dementia diagnoses. Participants underwent a series of longitudinal cognitive and psychometric assessments and gave written informed consent for the use of tissue samples and clinical information for research purposes. We selected samples from the BDR cohort for which DNA methylation data and genotype data were all available and for which all of the data passed quality control (QC), totaling 523 prefrontal cortex samples for subsequent validation analysis (**Supplementary Data 1, Table S1**). Details of DNA methylation and genetic data generation, processing, quality control and normalization from BDR cohort can be found in the original manuscript^50,51^.

Genotype data from BDR cohort were measured on the NeuroChip array which is a custom Illumina genotyping array with an extensive genome-wide backbone (n=306,670 variants) and custom content covering 179,467 variants specific to neurological diseases^52^. Genotype calling was performed using GenomeStudio (v2.0, Illumina) and quality control was completed using PLINK1.9^53^. The genetic data were then imputed by the Michigan Imputation Server^54^ (https://imputationserver.sph.umich.edu/index.html#!) which uses Eagle2^55^ to phase haplotypes, and Minimac4 (https://genome.sph.umich.edu/wiki/Minimac4) with the most recent 1000 Genomes reference panel (phase 3, version 5). Finally, imputed SNPs with minor allele frequency < 0.05, R^2^ INFO score < 0.5, Hardy-Weinberg equilibrium P-value < 10^−4^ and call rate < 0.95 were excluded. All samples with more than 5% missing were also excluded. The final quality controlled dataset of genotype consisted of 6,607,832 SNPs.

In short, DNA methylation data was generated on the dorsolateral prefrontal cortex [DLPFC] from each BDR donor. DNA was isolated from ∼100mg of tissue using the Qiagen AllPrep DNA/RNA 96 Kit (Qiagen, cat no.80311) following tissue disruption using BeadBug 1.5mm Zirconium beads (Sigma Aldrich, cat no. Z763799) in a 96-well Deep Well Plate (Fisher Scientific, cat no. 12194162) shaking at 2500 rpm for 5min. Genome-wide DNA methylation was profiled using the Illumina EPIC DNA methylation array (Illumina Inc.), which interrogates >850,000 DNA methylation sites across the genome^56^. After stringent data quality control using the *wateRmelon* package^57^ as previously described^50^, the BDR dataset consisted of DNA methylation estimates for 800,916 DNA methylation sites profiled.

### Genomic features annotation of c*PCDH* genes

c*PCDH* genes were annotated according to 3 genomic features. First, the CpGs (hg19 genome coordinate-*PCDHA*:140165721-140391929, *PCDHB*:140430979-140627802, *PCDHG*: 140710252-140892546) mapping to c*PCDH* loci were divided into three categories based on mean DNA methylation level: hypomethylated (β < 0.25), intermediately methylated (0.25 < β < 0.7) and hypermethylated (β > 0.7). The hypomethylated and hypermethylated thresholds were chosen based on the location of the hypo- and hyper-methylated peaks in the density plots of chromosome 5 methylation in BIOS discovery data (**Figure 1A**). Second, we annotated CpGs involved according to promoters related features: promoters and non-promoters. Transcripts database (genome hg19) were first downloaded from UCSC using *TxDb.Hsapiens.UCSC.hg19.knownGene* package, and then *promoters*() function for extracting the promoters information of every c*PCDH* members. Promoters were annotated as 1500 bp upstream of transcription start site (TSS) to 500 bp downstream of TSS, and all remaining regions as non-promoters. Finally, unique genomic regions of c*PCDH* were visualized with *Gviz* package^58^. To test whether hypomethylated CpGs in the c*PCDH* loci were enriched for promoter regions, we calculated odds ratios (OR) of c*PCDH* hypomethylated CpGs being annotated with promoters compared to all c*PCDH* non-hypomethylated CpGs being annotated as promoters using a Chi-square test.

**Figure 1.**
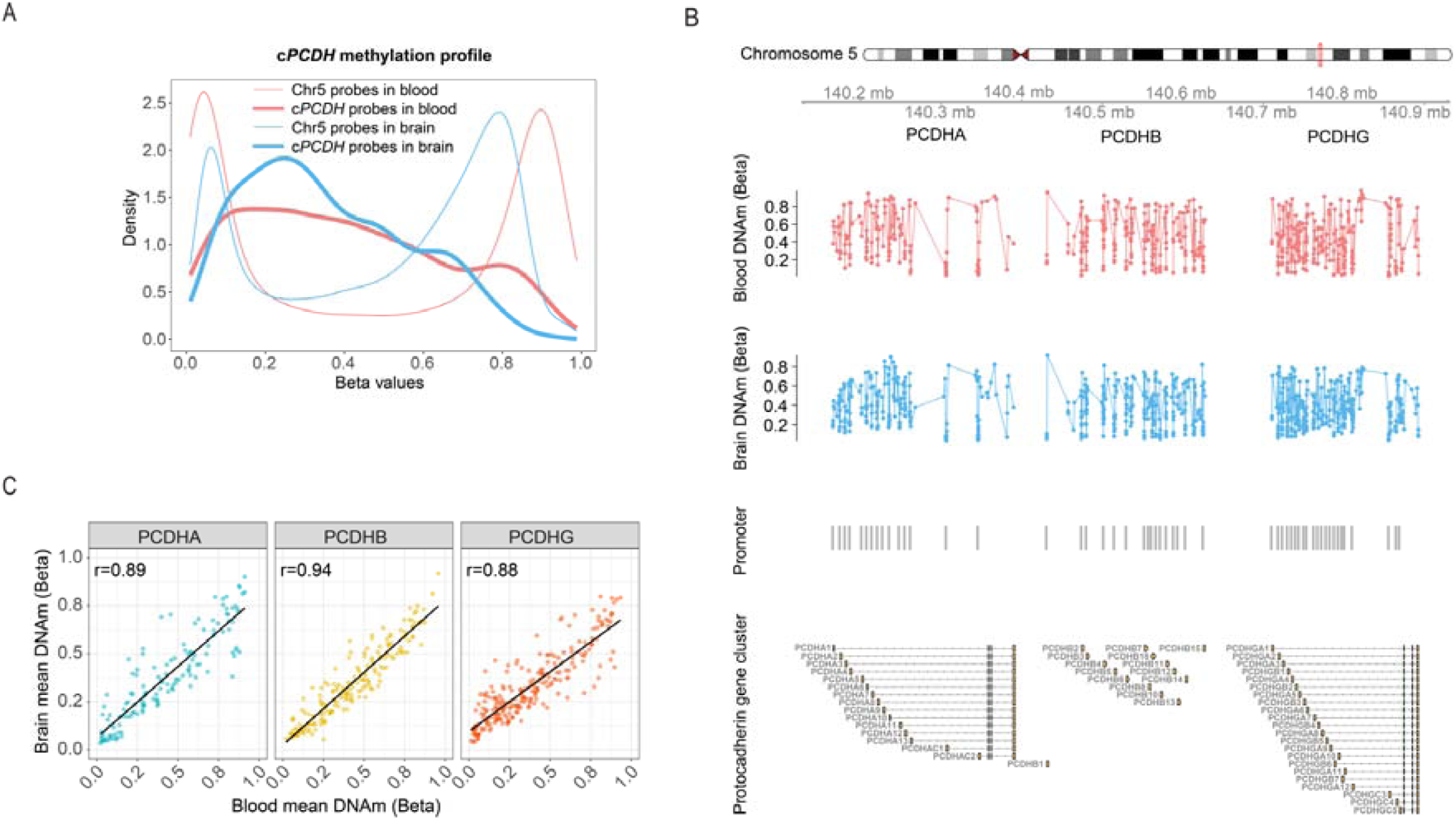
Overview of c*PCDH* methylation patterns in blood and brain (prefrontal cortex). **A** Distribution of c*PCDH* methylation based on 607 CpGs in BIOS blood (n=3777, Illumina 450k array) and based on 747 CpGs in BDR prefrontal cortex samples (n=523, Illumina EPIC array). The thicker red line and thicker blue line represent the distribution of c*PCDH* methylation in blood and brain (prefrontal cortex), respectively in contrast to chromosome 5 methylation distribution (thinner blue and red lines). **B** Four different tracks plotted by *Gviz* package^58^ to clearly display global methylation pattern in the c*PCDH* genes. First track shows methylation of 607 CpGs mapping to *PCDHA* (146 CpG), *PCDHB* (202 CpGs) and *PCDHG* (259 CpGs) in blood. Each red dot represents the mean methylation of one CpG in 3777 blood samples. Second track shows methylation of 747 CpGs mapping to *PCDHA* (199 CpG), *PCDHB* (240 CpGs) and *PCDHG* (308 CpGs) in brain (prefrontal cortex). Each blue dot represents the mean methylation of one CpG in 523 prefrontal cortex samples. Third track shows the location of promoter regions of each c*PCDH* member (1500 bp upstream of TSS to 500 bp downstream of TSS). Fourth track shows the genomic organization of the *PCDHA*, *PCDHB* and *PCDHG* gene clusters with exons. **C** Comparison of mean methylation level of overlapping c*PCDH* CpGs in blood and brain (prefrontal cortex). Pearson’s correlation coefficients are provided.

### Candidate gene analysis

Of the 9 genes previously reported to be involved in *cPCDH* regulation according to mouse studies^16,18,21–24,59,60^ and human cell lines^14,61^, 6 could be analyzed here. Three genes dropped out because no SNPs were available in blood (*MECP2*^61^) or brain (*WAPL*^23^ and *SETDB1*^16^). For the remaining genes, namely *SMCHD1*^21,22^, *CDCA7*^24^, *RAD21*^14^, *WIZ*^59^, *DNMT3B*^18^ and *CTCF*^14,60^, we fitted a linear regression model using R package *Matrix-eQTL*^62^ to test association between the SNPs near the 50bp window of candidate gene and 607 CpGs mapping to the c*PCDH* genes measured in whole blood of 3777 unrelated individuals using the Illumina 450k array, correcting for known covariates M (age, sex, cohort, cell counts and technical batches, namely sentrix position and sample plate) and 2 unknown hidden confounders U, estimated using *cate*^63^. Blood cell composition was estimated by R package *EpiDISH*^64^ and resulted in 12 broad immune cell type predictions including naïve CD4T cells, Basophils, memory CD4T cells, Memory B cells, Naive B cells, Regulatory T Cells, memory CD8 cells, naïve CD8T cells, Eosinophils, NK cells, Neutrophils and Monocytes. Neutrophils were excluded from the model to exclude collinearity so that the effect of this cell type becomes included in the intercept.

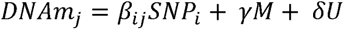

For each genetic variant SNP_i_, this model yields 607 P-values P_ij_. Next, we combined all 607 P-values corresponding to one genetic variant SNP_i_ into one overall *P*-value P_i_ using the *Simes* procedure for multiple testing correction^65^. SNPs with an overall *P*-value < 1 × 10^−5^ in candidate gene analysis were considered significantly associated with c*PCDH* methylation^66^. False discovery rate (FDR) was used to determine which of 607 CpGs were individually associated with the top SNP.

### Genome-wide of c*PCDH* methylation QTL analyses in blood

For genome-wide association analyses of c*PCDH* methylation, the same linear regression model was fitted as for the evaluation of candidate genes but now testing the association of 7,751,736 SNPs across the genome with each of 607 CpGs mapping to the c*PCDH* genes measured in whole blood of 3777 unrelated individuals. Likewise, we applied *Simes*^65^ procedure to perform multiple testing correction for 607 P-values corresponding to each genetic variant SNP_i_ and finally resulted in 7,751,736 overall P-value. This overall P-value per SNP indicates if an SNP influences DNA methylation anywhere on c*PCDH* genes. SNPs at genome-wide significance level of overall P-value < 5 × 10^−8^ were regarded to be associated with *PCDH* methylation. False discovery rate (FDR) was used to determine which of 607 individual CpGs were associated with the top SNP. Since we aimed to identify novel gene loci affecting c*PCDH* methylation *in trans*, SNPs in the c*PCDH* or mapping within 5MB of the cluster, defined as 5MB upstream of *PCDHA* transcription start sites (TSSs) to 5MB downstream of *PCDHG* transcription end sites (TESs), were removed from the analysis.

#### Local (*cis*) expression QTL mapping in blood

For *cis*-eQTL analysis in BIOS blood, we used a subset of 3031 individuals from the discovery cohort (N=3777, **Supplementary Data 1, Table S1**) for whom genetics, DNA methylation and also RNA-seq data were available. The association between the genotypes of a genetic variant SNP_i_ with the expression levels of genes Expression_j_, was tested by a linear model. Similar to the *trans*-meQTL mapping for c*PCDH* genes, we corrected for known covariates M (age, sex, cohort, cell counts and technical batches, namely flowcell number), and 2 unknown hidden confounders U using *cate*^63^.

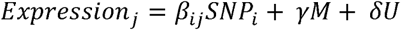

For this study, we only focus on top SNP in four genetic loci (*SMCHD1*, *OR2W3*, *SENP7* and *VENTX*) and tested its association with those genes that are identified as *cis*-eQTL according to eQTLGen consortium^67^. The Bonferroni correction was performed to the corresponding P-values *P*_ij_ to identify genes associated with the genetic variant SNP_i_ .

#### Validation of *trans*-meQTL in brain

To evaluate the effect of top SNP derived from blood in genetic loci on c*PCDH* methylation in brain, we repeated methylation QTL analysis with following statistical model using BDR cohort.

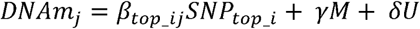

where DNAm_j_ represent 747 c*PCDH* methylation sites (EPIC array) matrix. M represent age, sex, experimental batch and derived brain cell proportions. Cell proportion estimates were derived from bulk cortex DNA methylation data using the Houseman method^68^, implemented with *minfi* functions and default parameters, incorporating the novel reference DNA methylation data generated on three Fluorescence-activated nuclei sorting (FANS)-purified nuclei populations (NeuN+, SOX10+, and NeuN–/SOX10–) from 12 DLPFC samples^50^. U represent PC1 which accounted for residual structure in the data. In brain, we tested top SNP from 6 candidate genetic loci and 3 additional genetic loci as detected in blood. For top SNP SNP_top_i_ in each genetic locus (i = S*MCHD1*, *CDCA7*, *RAD21*, *WIZ*, *DNMT3B*, *CTCF*, *OR2W3*, *SENP7* and *VENTX*), this model yields 747 P-values P_j_ . The validation in the brain of results in blood was assessed by two aspects: (1) The statistical consistency for detected significant CpGs in blood and brain. To test the probability of statistically significant CpGs in blood also being significant in brain, we calculated odds ratio (OR) of CpGs significant in blood to be also significant in brain. Statistical significance was evaluated using a Chi-square test or a Fisher’s exact test (if expected cell counts were < 5). (2) The consistency of effect direction for detected significant CpGs in blood or brain. Summarizing the number of CpGs significant in blood or brain with same effect direction (Positive + Negative), which included three categories of CpGs: Blood&Brain (statistically significant both in blood and brain), Blood only (only FDR significant in blood) and Brain only (only nominally significant in brain). Binomial test was used to determine whether direction of effects were concordant.

## Results

### c*PCDH* DNA methylation patterns are similar in blood and brain

In whole blood, we evaluated methylation levels of 607 CpGs (Illumina 450k array) mapping to the c*PCDH* genes using a data set of 3777 samples (**Supplementary Data 1, Table S2**). In contrast to the typical bimodal distribution of genomic DNA methylation with the majority of CpGs being either hypo- or hypermethylated, the majority of CpGs in c*PCDH* genes showed intermediate DNA methylation levels (**Figure 1A**). We found a similar result in brain (**Figure 1A**), where we evaluated the methylation levels of 747 CpGs (Illumina EPIC array) located in the c*PCDH* genes using a data set of 523 prefrontal cortex samples (**Supplementary Data 1, Table S3**).

To gain more insight in the pattern of c*PCDH* methylation, we first examined the mean methylation level for each CpG mapping to the *PCDHA*, *PCDHB* and *PCDHG* gene subclusters in blood and brain (**Figure 1B**). As shown in **Figure 1B**, the *PCDHA* and *PCDHG* subclusters have a similar genomic architecture where tandem arrays of alternative exons are followed by constant exons, while the *PCDHB* subcluster contains single exon genes. We observed that c*PCDH* methylation levels varied over short genomic distances between hypo-, intermediate and hypermethylation where hypomethylation coincided with promotor regions. In both blood and brain, promoter CpGs were more likely to be hypomethylated as compared with non-promoter CpGs (OR_blood_ = 6.0, P=1.1×10^-20^; OR_brain_=4.4, P=7.9×10^-18^; **Supplementary Data 1, Table S4**). Finally, we tested the correlation between DNA methylation in blood and brain for the 562 overlapping CpGs. The correlation was high ranging between 0.88 for the *PCDHG* subcluster to 0.94 for the *PCHDB* subcluster (**Figure 1C**).

### The effect of candidate gene loci on c*PCDH* methylation in blood and brain

The candidate genes *SMCHD1*^21,22^, *CDCA7*^24^, *RAD21*^14^, *WIZ*^59^, *DNMT3B*^18^ and *CTCF*^14,60^ were evaluated for an effect on c*PCDH* methylation in human blood and brain (**Figure 2A**). We found that top SNP *rs10775431* at the *SMCHD1* locus was significantly associated with c*PCDH* methylation in blood (P_Simes_ =2.7×10^-6^, minimal nominal P= 4.4×10^-9^) and affected 221 c*PCDH* CpGs methylation in blood (P_FDR_<0.05, 450k array, **Supplementary Data 1, Table S5; Supplementary Data 2, Table S6**). Of those, 205 were also measured in brain (EPIC array) and 66 (32%) were validated in brain (P<0.05; OR = 3.0, P=1.0×10^-7^; **Figure 2B, Supplementary Data 2, Table S6-S7, Table S18**). In addition, among the 253 CpGs (available from both the 450k and EPIC array) found to be associated with the top SNP in either blood or brain, 230 (91%) had a consistent direction of effect (P=1.0x10^-8^, **Supplementary Data 2, Table S6-S7**). Interestingly, c*PCDH* CpGs affected by the *SMCHD1* variant in blood or brain occurred across the whole cluster and were both hypo-, intermediate and hypermethylated (**Figure 2C**).

**Figure 2.**
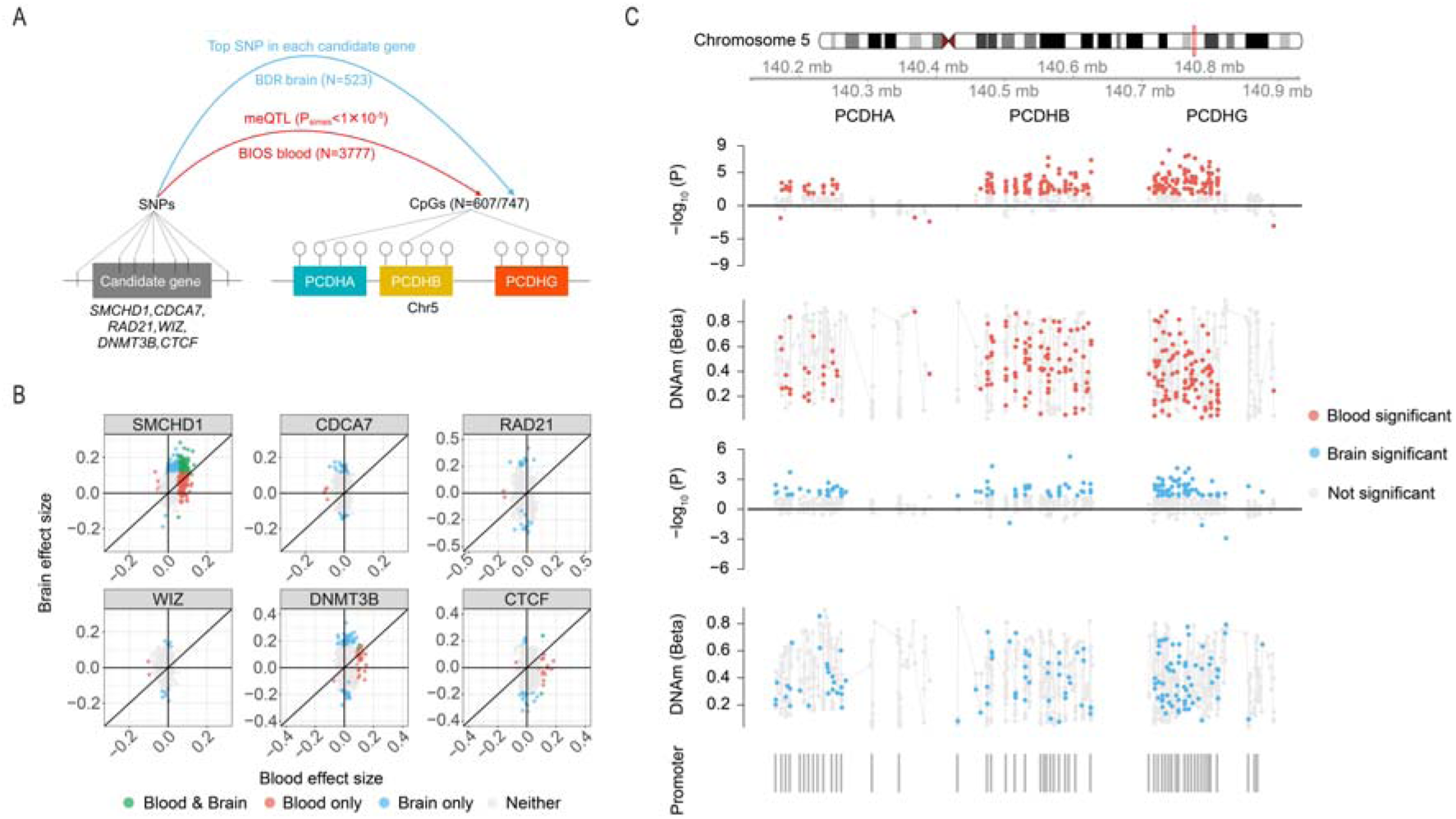
Effect of 6 candidate gene loci on c*PCDH* methylation in blood and brain (prefrontal cortex). **A** We first performed *trans*-meQTL analysis to test the association between the SNPs near the 50bp window of candidate gene and 607 c*PCDH* CpGs in BIOS blood samples (n=3777). SNPs with an overall P-value < 1 × 10^−5^ in candidate gene analysis were considered significantly associated with c*PCDH* methylation. Then the effect of top SNP in each candidate gene was validated in BDR prefrontal cortex samples (n=523). **B** Scatter plot of effect sizes for top SNP in 6 candidate genes on c*PCDH* methylation observed in blood and brain (prefrontal cortex). Green dots represent CpGs statistically significant in both blood and brain (prefrontal cortex). Red and blue dots represent CpGs statistically significant only in blood or brain (prefrontal cortex), respectively. Gray dots represent CpGs statistically significant in neither blood nor brain (prefrontal cortex). **C** Effect of *SMCHD1* variant (*rs10775431*) on c*PCDH* methylation in blood and brain (prefrontal cortex). The tracks display the P-values resulting from the statistical test, the direction of effect and mean methylation level of c*PCDH* CpGs affected by *SMCHD1* variant (*rs10775431*), respectively for blood and brain (prefrontal cortex). Red and blue dots represent CpGs that were statistically significant in blood (P_FDR_<0.05) and brain (nominal *P*-value<0.05, prefrontal cortex), respectively. The fifth track shows the location of promoter regions of each c*PCDH* member (1500 bp upstream of TSS to 500 bp downstream of TSS).

Further indicating that the genetic associations were mediated by *SMCHD1*, the top SNP *rs10775431* was a blood expression QTL of the gene according to BIOS (**Supplementary Data 3, Table S19**) and eQTLGen consortium^67^. Further, for *DNMT3B* and *CTCF* we only observed weak evidence for consistent associations with c*PCDH* methylation in blood and brain, while top SNP at the remaining candidate genes was not associated as none of CpGs that were statistically significant in blood validated in brain (**Supplementary Data 2, Table S8-S18**). Overall, genetic variation at the *SMCHD1* locus showed by far the most strongest and most consistent effect on c*PCDH* methylation in blood and brain of the 6 candidate loci evaluated.

#### Genome-wide identification of genetic loci affecting c*PCDH* methylation in blood

Next, we aimed to identify new loci *in trans* affecting c*PCDH* methylation. In view of the consistent DNA methylation patterns and effects of candidate genes in blood and brain, we set out to perform a genome-wide association analysis in blood, the data set with the largest sample size (N=3777) (**Figure 3A**). We detected 334 genetic variants representing 7 independent loci associated with c*PCDH* methylation *in trans* affecting at least 1 out of the 607 tested CpGs (P_Simes_ <5×10^-8^, **Figure 3B**, **Supplementary Data 1, Table S5**) thus excluding *cis*-eQTL effects defined as within 5MB of c*PCDH* genes. Analysis of the top SNPs per locus showed that 4 out of 7 loci were associated with 1 c*PCDH* CpG only (P_FDR_<0.05; **Table 1**). We did not further examine these loci. The 3 remaining top SNPs were located on chromosomes 1, 3 and 10, and were associated with widespread c*PCDH* methylation *in trans*, each affecting at least 50 c*PCDH* CpGs (P_FDR_<0.05; **Table 1**). Further inspection of all suggestive genome-wide associations (P_Simes_<1×10^-5^), excluded the possibility that we missed loci with widespread effects similar to *SMCHD1* (**Supplementary Data 1, Table S5**).

**Figure 3.**
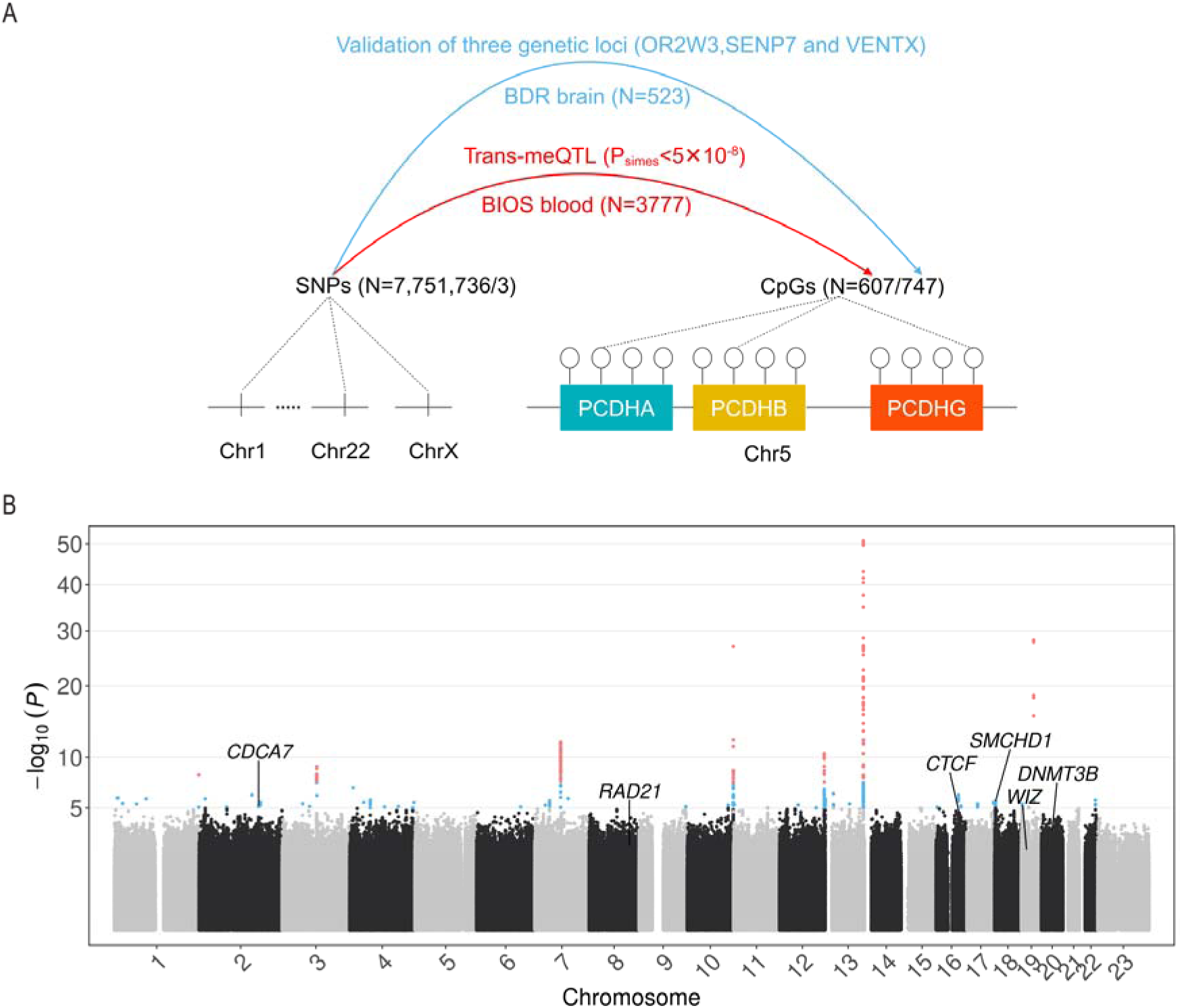
Genome-wide of c*PCDH* methylation QTL analyses. **A** To identify additional novel genetic loci influencing c*PCDH* methylation, we performed *trans*-meQTL analysis to test the association between the SNPs across whole genome and 607 c*PCDH* CpGs in BIOS blood (n=3777). Then the effect of three novel genetic loci were validated in BDR prefrontal cortex samples (n=523). **B** Manhattan plot showing all tested *trans*-SNPs for an overall effect on c*PCDH* methylation in blood. Significant associations are depicted in red (P_Simes_ <5×10^-8^) and blue dots indicate SNPs with overall P-values between 1×10^-5^ (suggestive threshold) and 5×10^-8^ (genome-wide threshold). Additionally, italic text represents the location of 6 candidate gene loci (top SNP in candidate gene analysis) known to be involved in c*PCDH* regulation.

**Table 1.**
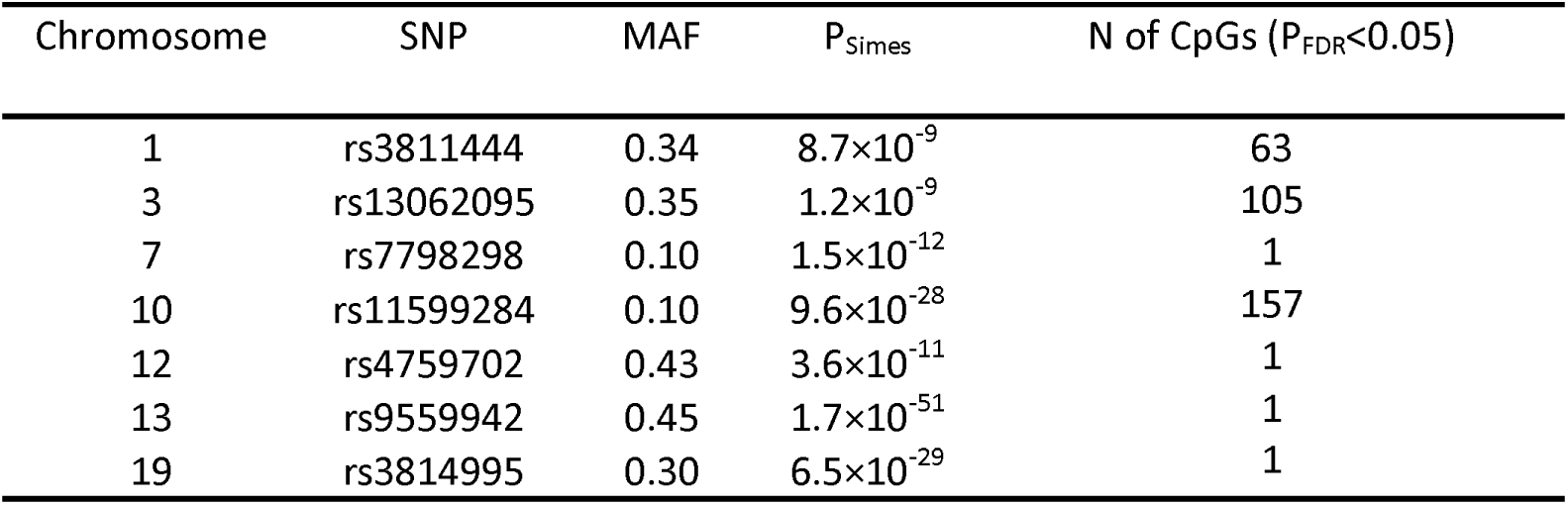
Effect of seven top SNPs on *PCDH* methylation in blood.

### No validation of the *OR2W3 locus*effect in brain

The top SNP *rs3811444* is a missense variant (MAF=0.34) on chromosome 1, located in *TRIM58* (**Figure 4A**). By looking up eQTLGen consortium^67^ (n=26395), we found that *rs3811444* in blood only regulates nearby *OR2W3*, and its T allele was associated with increased expression. This was also the case in BIOS blood data (P=2.1×10^-10^; **Supplementary Data 3, Table S19**).

**Figure 4.**
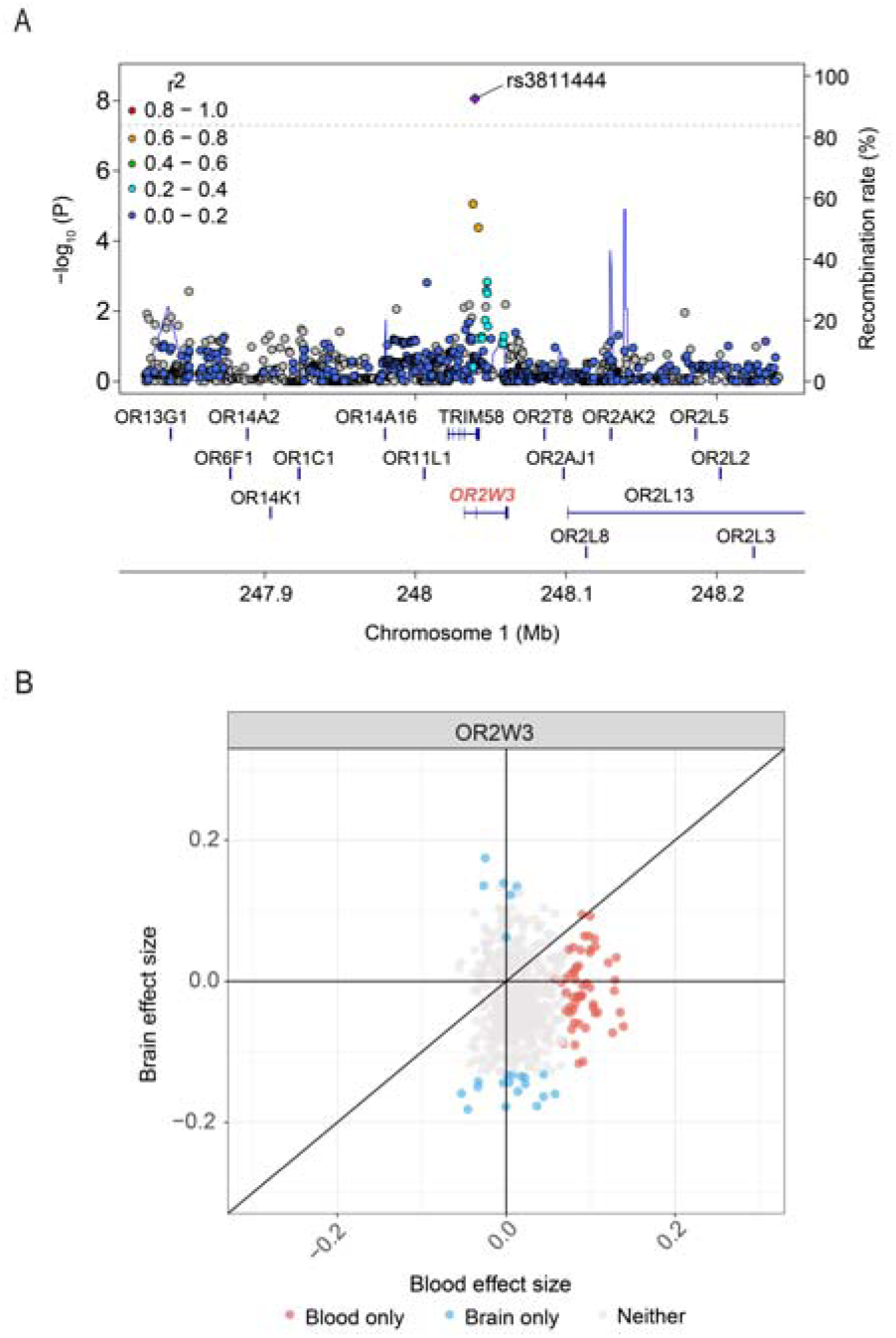
Effect of *OR2W3* locus on c*PCDH* methylation in blood and brain (prefrontal cortex). **A** Locus Zoom plot of top SNP *rs3811444* on chromosome 1. **B** Scatter plot of effect sizes for top SNP *rs3811444* in the *OR2W3* genetic locus on c*PCDH* methylation observed in blood and brain (prefrontal cortex). Red and blue dots represent CpGs statistically significant only in blood or brain (prefrontal cortex), respectively. Gray dots represent CpGs statistically significant in neither blood nor brain (prefrontal cortex).

The effect of *rs3811444* on c*PCDH* methylation observed in blood, however, was absent in brain samples. First, we found *rs3811444* affected 63 *PCDH* CpGs methylation in blood (450k array, **Table 1**). Of those, 55 were also measured in brain (EPIC array) and none of the statistically significant CpGs in blood were validated in brain (**Supplementary Data 3, Table S20-21; Table S26**). Second, among the 78 CpGs (available from both the 450k and EPIC array) found to be associated with the top SNP in either blood or brain, only 30 CpGs (38%) detected significant in blood or brain had a consistent direction of effect (**Figure 4B; Supplementary Data 3, Table S20-21**).

### Validating the *SENP7 locus* effect in brain

The top SNP *rs13062095* (MAF=0.35) in the second locus is located between *SENP7* gene and *TRMT10C* gene on chromosome 3, which marked a region with SNPs in strong linkage disequilibrium (r^2^>0.9) overlapping with the *SENP7* gene (**Figure 5A**). According to the eQTLGen consortium^67^ (n=26,395), the C allele of *rs13062095* was associated with the expression of several nearby genes including *SENP7, IMPG2*, *CEP97*, *TFG* and *NXPE3*, but most strongly with the expression of the *SENP7* gene. In BIOS blood, we found that all genes except for *IMPG2* were also significantly associated with the C allele of *rs13062095* (**Supplementary Data 3, Table S19**). Interestingly, the three genes, *SENP7*, *IMPG2* and *CEP97*, were also *cis*-eQTL gene of *rs13062095* in cortex (MetaBrain consortium^69^, n=2757). In cortex, the SNP was associated with the expression of five genes in *trans* (MetaBrain consortium^69^, n=2757), four of which were protocadherin genes (*PCDHB5*, *PCDHGA1*, *PCDHGB1* and *PCDHGB5*) and one zinc finger coding gene (*ZNF274*), which has recently been implicated in c*PCDH* gene regulation^70^. However, genetic variation at *ZNF274* was not associated with c*PCDH* methylation in our study (P_simes_>10^-5^) and this was also case in GoDMC^71^ and Pan-mQTL^72^.

**Figure 5.**
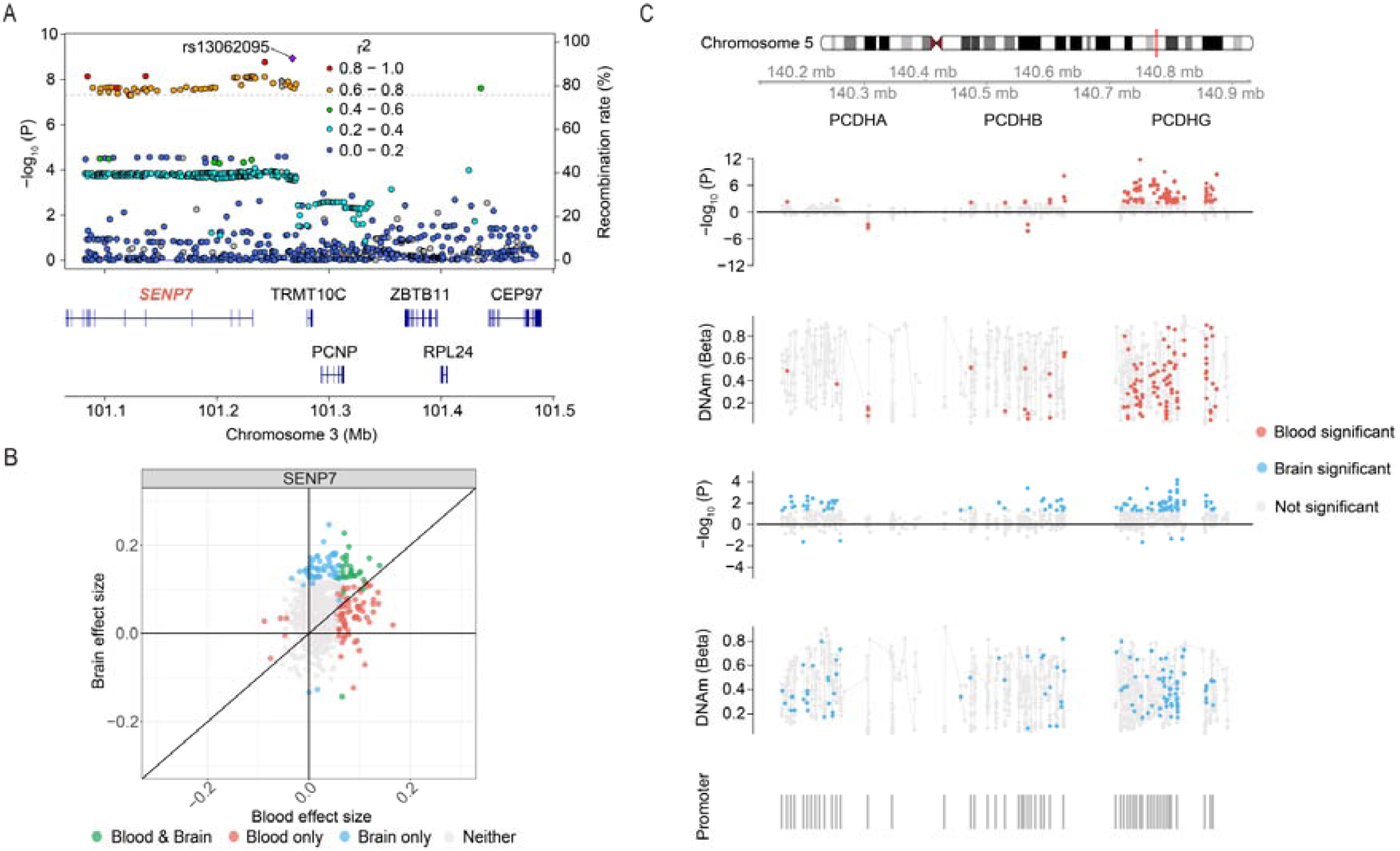
Effect of *SENP7* locus on c*PCDH* methylation in blood and brain (prefrontal cortex). **A** Locus Zoom plot of top SNP *rs13062095* on chromosome 3. **B** Scatter plot of effect sizes for top SNP *rs13062095* in *SENP7* genetic locus on c*PCDH* methylation observed in blood and brain (prefrontal cortex). Green dots represent CpGs statistically significant in both blood and brain (prefrontal cortex). Red and blue dots represent CpGs statistically significant only in blood or brain (prefrontal cortex), respectively. Gray dots represent CpGs statistically significant in neither blood nor brain (prefrontal cortex). **C** Effect of *SENP7* variant (*rs13062095*) on c*PCDH* methylation in blood and brain (prefrontal cortex). The tracks display the P-values resulting from the statistical test, the direction of effect and mean methylation level of c*PCDH* CpGs affected by *SENP7* variant (*rs13062095*), respectively for blood and brain (prefrontal cortex). Red and blue dots represent CpGs that were statistically significant in blood (P_FDR_<0.05) and brain (nominal P-value<0.05, prefrontal cortex), respectively. The fifth track shows the location of promoter regions of each c*PCDH* member (1500 bp upstream of TSS to 500 bp downstream of TSS).

Next, we tested the association of the top SNP *rs13062095* on c*PCDH* methylation in blood, and then validated in brain. A high concordance was observed between the two tissues (**Figure 5B, Supplementary Data 3, Table S22-23; Table S26**). First, top SNP *rs13062095* affected methylation of 105 c*PCDH* CpGs in blood (P_Simes_ =1.2×10^-9^, minimal nominal P= 1.9×10^-12^, 450k array, **Table 1**). Of these, 97 were also measured in brain (EPIC array) and 26 (27%) were validated in brain (P<0.05; OR = 2.6, P=2.4×10^-4^). Second, among the 154 CpGs (available from both the 450k and EPIC array) found to be associated with the top SNP in either blood or brain, 130 CpGs (84%) detected as significant in blood or brain had consistent direction of effect (P=10^-8^).

The high *SENP7* expression associated C-allele of *rs13062095* was associated with overall higher c*PCDH* methylation. Inspection of the three subclusters revealed this effect was mainly apparent for *PCDHG* in blood but had an effect on all three subclusters in brain and concerned hypo-, intermediate and hypermethylated CpGs (**Figure 5C**). To check concordance, we further looked up *rs13062095* in large-scale blood meQTL consortia datasets including GoDMC^71^ and Pan-mQTL^72^. In GoDMC^71^, *rs62280667*, a SNP in high LD with *rs13062095* (R^2^=0.97), was associated with one c*PCDH* CpG methylation (**Supplementary Data 3, Table S27**). In Pan-mQTL^72^, we found that *rs13062095* itself was associated with the methylation of three c*PCDH* CpGs in *trans* (**Supplementary Data 3, Table S28**).

### Validating the *VENTX locus* effect in brain

The top SNP *rs11599284* (MAF=0.1) of the third identified locus is a missense variant on chromosome 10 (glycine to arginine substitution at amino acid 73), located in the *UTF1* gene (**Figure 6A**). Moreover, we found that the A allele of *rs11599284* was associated with the expression of several nearby genes including *VENTX E, CHS1*, *TUBGCP2* and *ADAM8*, but most strongly with the expression of *VENTX* (eQTLGen consortium^67^, n=26,395). In BIOS blood data (n=3777), the strong *cis*- effect for *VENTX* was not observed, while the *TUBGCP2* and *ADAM8* were consistently found (**Supplementary Data 3, Table S19**). The SNP was not evaluated in the MetaBrain Consortium^69^, precluding an assessment whether it was a *cis*-eQTL in brain.

**Figure 6.**
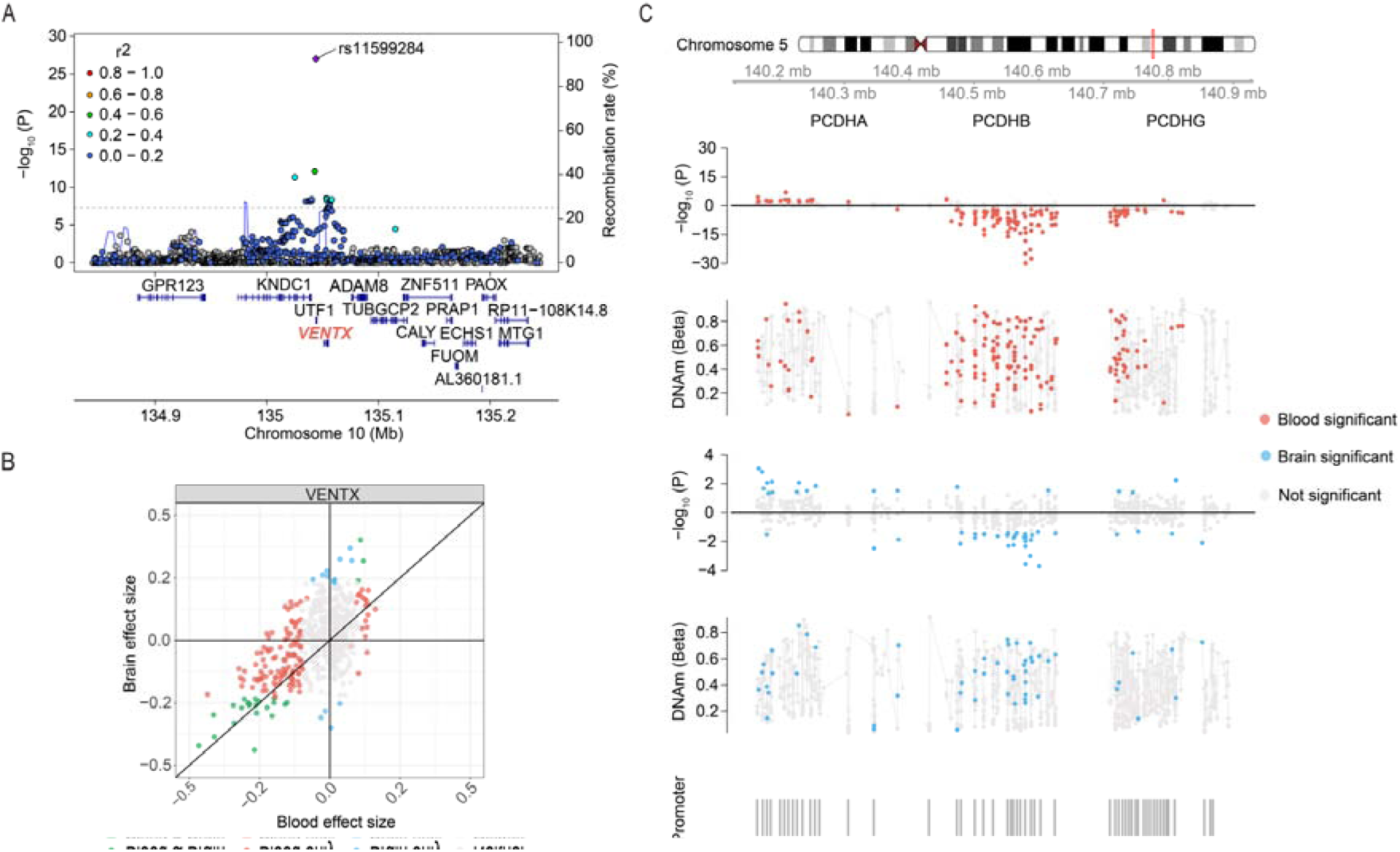
Effect of *VENTX* locus on c*PCDH* methylation in blood and brain (prefrontal cortex). **A** Locus Zoom plot of top SNP *rs11599284* on chromosome 10. **B** Scatter plot of effect sizes for top SNP *rs11599284* in *VENTX* genetic locus on c*PCDH* methylation observed in blood and brain (prefrontal cortex). Green dots represent CpGs statistically significant in both blood and brain (prefrontal cortex). Red and blue dots represent CpGs statistically significant in only blood or brain (prefrontal cortex), respectively. Gray dots represent CpGs statistically significant in neither blood nor brain (prefrontal cortex). **C** Effect of *VENTX* variant (*rs11599284*) on c*PCDH* methylation in blood and brain (prefrontal cortex). The tracks display the P-values resulting from the statistical test, the direction of effect and mean methylation level of c*PCDH* CpGs affected by *VENTX* variant (rs11599284), respectively for blood and brain (prefrontal cortex). Red and blue dots represent CpGs that were statistically significant in blood (P_FDR_<0.05) and brain (nominal P-value<0.05, prefrontal cortex), respectively. The fifth track shows the location of promoter regions of each c*PCDH* member (1500 bp upstream of TSS to 500 bp downstream of TSS).

Next, we tested the effect of the top SNP *rs11599284* on c*PCDH* methylation in blood, and then validated in brain. A high concordance was observed between the two tissues (**Figure 6B, Supplementary Data 3, Table S24-26**). First, top SNP *rs11599284* affected methylation of 157 c*PCDH* CpGs in blood (P_Simes_ =9.6×10^-28^, minimal nominal P= 1.6×10^-30^,450k array, **Table 1**). Of those, 150 were also measured in brain (EPIC array) and 24 (16%) were validated in brain (P<0.05; OR = 5.4, P=1.4×10^-7^). Second, among the 164 CpGs (available from both the 450k and EPIC array) found to be associated with the top SNP in either blood or brain, 130 CpGs (79%) detected as significant in blood or brain had a consistent direction of effect (P=10^-8^).

The A-allele of *rs11599284*, leading to the amino-acid change in *UTF1* and associated with low *VENTX* expression, was associated with overall lower c*PCDH* methylation. Inspection of the three subclusters revealed that this effect was most apparent for *PCDHB* and concerned hypo-, intermediate and hypermethylated CpGs (**Figure 6C**). To check concordance, we further looked up *rs11599284* in large-scale blood meQTL consortium include GoDMC^71^ and Pan-mQTL^72^. In GoDMC^71^, the SNP was not evaluated. In Pan-mQTL^72^, we found that this SNP was associated with the methylation of seven c*PCDH* CpGs in *trans* and all were mapping to the *PCDHB* subcluster (**Supplementary Data 3, Table S29**).

## Discussion

Our genetic study identified loci with a widespread effect on c*PCDH* methylation in human blood and brain. In contrast to many other genomic regions^73,74^, c*PCDH* methylation was highly correlated between blood and prefrontal cortex and the methylation patterns were concordant, which was characterized by a variation between hypomethylation near promoter regions to intermediate and hypermethylation in-between promoters. In a candidate gene analysis, we found that genetic variation at the *SMCHD1* gene affected methylation at all three *PCDH* subclusters in blood and brain, while no or a much less pronounced effects were observed for other candidate genes evaluated. In a genome-wide analysis, we identified two novel genetic loci, *SENP7* and *UTF1*/*VENTX*. The *SENP7* locus was primarily associated with *PCDHG* methylation in blood but had an effect on all three subclusters in brain. The *UTF1*/*VENTX* locus mainly influenced methylation of the *PCDHB* subcluster in blood and brain, and had less pronounced effects on the other subclusters. Our systematic genome-wide analysis identified novel genetic loci for c*PCDH* methylation regulation, significantly expands upon genes that were previously implicated in association with c*PCDH* methylation using mouse models, and identifies the direction of the effect on c*PCDH* methylation.

While previous work in mice established the critical role for multiple genes in c*PCDH* regulation^18,21,22,24,59,60^, their effects in c*PCDH* methylation in humans are only partly understood^12,13^. Here, we confirm the role of the *SMCHD1* locus in c*PCDH* methylation in human blood and prefrontal cortex and highlight its broad impact on methylation of all three *PCDH* subclusters. We also observed that other candidate gene loci had no or a much less pronounced effects on c*PCDH* methylation. For example, the *DNMT3B* locus showed weak evidence for consistent associations with c*PCDH* methylation in blood and brain. Given the known role of *DNMT3B* in *de novo* methylation, this may be explained by the minor effect of common genetic variation on *DNMT3B* expression. Studies with a larger number of blood and brain samples will likely be able to detect more associations due to a higher statistical power. The *CDCA7* locus is another candidate gene for which we did not detect consistent associations with c*PCDH* methylation in blood and brain. We previously reported that genetically controlled differential expression of *CDCA7* induces genome-wide DNA methylation changes in particular at repeats in humans indicating that common genetic variation does affect DNA methylation^45^. In addition, *CDCA7* plays a key role in c*PCDH* regulation in mouse neurons^24^. These apparent discrepancies may arise from either brain-specific effects which we could detect less well due to the lower number of brains samples in our analysis or from design differences between experimental study (tissues/cells from patients and knock-out/knock-in models for diseases) and our study (common genetic variation in general population).

Our genome-wide association study implicates the *SENP7* locus in c*PCDH* methylation regulation. *SENP7* is a strong candidate to influence c*PCDH* methylation based on following evidence. First, it was reported to affect methylation levels of multiple CpG sites in *trans*^45^. Second, *SENP7* has been shown to be involved in the deSUMOylation of the chromatin repressor *KAP1*/*TRIM28*^25^, which functions in DNA repair and epigenetic regulation including DNA methylation regulation^75,76^. Therefore, we hypothesize that *SENP7* affects c*PCDH* methylation through its interaction with *KAP1*. Third, using external large-scale methylation or expression QTL atlas including GoDMC^71^, Pan-mQTL^72^ and MetaBrain^69^, we observe that *SENP7* variant does affect c*PCDH* methylation and regulate c*PCDH* expression. Of note, when comparing our results with other published large-scale blood meQTL results, it is important to be aware of the differences in study design. The number of CpGs evaluated in genome-wide trans-meQTL analyses is substantially higher than in our study (607 c*PCDH* CpGs only) and hence, significant threshold cutoff for SNP-CpG pairs is several order of magnitude lower than in our analysis focused on c*PCDH*. Interestingly, c*PCDH* differential methylation has been associated with many neurological diseases^3–6^ and one GWAS study indicated that the *SENP7* locus associated with late-onset Alzheimer’s disease^77^. Another study found that c*PCDH* methylation alterations related to a decline in regional brain volume and cognitive skills^78^. It will be interesting to establish in future studies whether *SENP7* is involved in Alzheimer’s disease through its effect on c*PCDH* methylation.

Similarly, our genome-wide analysis indicated a role for the *UTF1*/*VENTX* locus in c*PCDH* methylation regulation. These genes have previously not been implicated in epigenetic regulation. *UTF1* is required for the proper differentiation of embryonic stem cells^26^ while *VENTX* regulates gene expression in stem cells and during embryonic development as a transcription factor^27^. The SNP we identified, *rs11599284*, is associated with the expression of *VENTX* and a missense variant resulting in a glycine to arginine substitution in the *UTF1* protein^79^. The missense mutation is predicted to be deleterious, while experimental evidence implicated the mutation in decrease in chromatin association, albeit only in combination with a second mutation^79^.

Our study contributes to a better understanding of the epigenetic regulation of the c*PCDH* genes through identification of *SENP7* and *UTF1*/*VENTX* as new loci influencing c*PCDH* methylation across blood and brain. However, our study also has limitations. First, we first analyzed blood samples followed by validation in brain, which means that we missed brain-specific genetic loci influencing c*PCDH* methylation. Second, our brain sample size is relatively small (N=523) precluding a genome-wide analysis to capture loci influencing c*PCDH* methylation in brain. Third, our results does not provide insight into the exact underlying mechanisms of *SENP7* and *UTF1*/*VENTX* function in c*PCDH* regulation. In particular for the functionally poorly characterized *UTF1* and *VENTX* genes it is currently unclear how they could influence epigenetic regulation. Functional experiments will be required to demonstrate causality for the genes we implicated in c*PCDH* methylation in blood and brain and uncover the mechanisms involved.

In conclusion, we identified two new loci associated with c*PCDH* methylation across blood and brain on top of *SMCHD1*, namely *SENP7* and *UTF1*/*VENTX*. These results can guide follow-up studies into the epigenetic regulation of c*PCDH* genes, the role of c*PCDH* in human traits and diseases, and may eventually lead to a better understanding of the generation of neuronal cell surface barcodes.

## Availability of data and materials

The BIOS whole blood data are available from the European Genome-Phenome Archive (EGAC00001000277). The BDR brain data are available from the Dementias Platform UK (DPUK) (https://portal.dementiasplatform.uk/CohortDirectory/Item?fingerPrintID=BDR) upon application. Scripts for the main analyses in this manuscript are available at GitHub [https://github.com/YunfengLUMC/cPCDH_methylation_regulation] under an opensource MIT license. The summary blood meQTL results from the Europe population (GoDMC) are available at http://mqtldb.godmc.org.uk. The summary blood meQTL results from the Asian population are available at https://www.biosino.org/panmqtl/home. The summary blood eQTL results (eQTLGen consortium) are available at https://www.eqtlgen.org. The summary brain eQTL results (MetaBrain consortium) are available at https://www.metabrain.nl.

## Acknowledgements

BIOS Consortium samples and data were contributed by LifeLines, the Leiden Longevity Study, the Netherlands Twin Registry (NTR), the Rotterdam Study, the Genetic Research in Isolated Populations program, the Cohort on Diabetes and Atherosclerosis Maastricht (CODAM) study, and the Prospective ALS study Netherlands (PAN). The BIOS was funded by BBMRI-NL (a Research Infrastructure financed by the Dutch government (NWO, numbers 184.021.007 and 184.033.111)). Cortex DNA methylation data generation was funded by Medical Research Council (MRC) grants K013807 and W004984 (awarded to J.M.). Data analysis was undertaken using high-performance computing supported by a Medical Research Council (MRC) Clinical Infrastructure award (M008924 awarded to J.M.). The BDR is jointly funded by Alzheimer’s Research UK (ARUK) and the Alzheimer’s Society in association with the Medical Research Council (MRC). LD was supported by NWO-VIDI grant (ZonMW91718350). YL was supported by a PhD fellowship from the China Scholarship Council (CSC). We thank the participants of all aforementioned biobanks and acknowledge the contributions of the investigators to this study. This work was carried out on the Dutch national e-infrastructure with the support of SURF Cooperative.

## Authors’ contributions

Conceptualization: BTH, LD and JM. Methodology: YL, BTH. Formal analysis: YL, EH and MV. Resources: BIOS Consortium and BDR consortium. Writing – original draft: YL, BTH. Writing – review and editing: BTH, MV, EH, HM, JM and LD. Visualization: YL, BTH. Supervision: BTH and LD. All authors have read and approved the final manuscript.

## Declaration of interests

The authors declare that they have no competing interests.

## Ethics declarations

### Ethics approval and consent to participate

The study was approved by the institutional review boards of the participating centers (CODAM, Medical Ethical Committee of the Maastricht University; LL, Ethics Committee of the University Medical Centre Groningen; LLS, Ethical Committee of the Leiden University Medical Center; PAN, Institutional Review Board of the University Medical Centre Utrecht; NTR, Central Ethics Committee on Research Involving Human Subjects of the VU University Medical Centre; RS, Institutional Review Board (Medical Ethics Committee) of the Erasmus Medical Center; BDR, Ethical Committee of University of Exeter Medical School Research). All participants have given written informed consent, and the experimental methods comply with the Helsinki Declaration.

## Supplementary Information

*Supplementary Data 1*.

Contains **Table S1-S5**.

*Supplementary Data 2*.

Contains **Table S6-S18**.

*Supplementary Data 3*.

Contains **Table S19-29**.

## Notes

### Competing Interest Statement

The authors have declared no competing interest.

